# Induced mineralization in *Escherichia coli* biofilms: the key role of bacterial alkaline phosphatase

**DOI:** 10.1101/2022.09.29.509673

**Authors:** Laura Zorzetto, Ernesto Scoppola, Emeline Raguin, Kerstin G. Blank, Peter Fratzl, Cécile M. Bidan

## Abstract

Biofilms appear when bacteria colonize a surface and synthesize and assemble extracellular matrix components. In addition to the organic matrix, some biofilms precipitate mineral particles such as calcium phosphate. While calcified biofilms induce diseases like periodontitis in physiological environments, they also inspire the engineering of living composites. Understanding mineralization mechanisms in biofilms will thus provide key knowledge for either inhibiting or promoting mineralization in these research fields. The enzyme alkaline phosphatase (ALP) plays a key role in calcium phosphate precipitation in mammalian bone tissue. Produced by eukaryotic cells, ALP catalyzes the hydrolysis of monophosphates starting from different precursors (e.g., alkaloids, proteins) and makes phosphate ions readily available for the precipitation with calcium. Bacterial ALPs are expressed by the well-characterized gram-negative and gram-positive bacteria *E. coli* and *S. aureus* as well as a large number of marine and soil bacteria. While it was recently proposed that bacterial ALPs induce mineral precipitation, their role in biofilm mineralization is not fully understood. In this work, we address this question using the biofilm-forming *E. coli* K-12 strain W3110, which expresses periplasmic ALP from the *phoA* gene. We first identify the mineralization conditions of biofilms grown on nutritive agar substrates supplemented with calcium ions and β-glycerophosphate. We then localize the mineral phase at different scales, using light and scanning electron microscopy as well as X-ray microtomography. Wide-angle X-ray scattering enables us to further identify the mineral as being hydroxyapatite. Finally, growing *E. coli* cells on mineralizing medium supplemented with an ALP inhibitor demonstrates that ALP is essential for biofilm mineralization. This is confirmed with a bacteria-free model, where the deposition of a drop of bacterial ALP solution on calcium and β-glycerophosphate containing agar substrate is sufficient to induce mineralization. Overall, these results will benefit the development of strategies against diseases involving calcified biofilms as well as the engineering of biofilm-based living composites.

## Introduction

Microbial biofilms form when bacteria colonize a surface and synthesize extracellular matrix components to survive challenging environments.^1^ In addition to the organic matrix, some biofilms accumulate mineral particles.^2^ Microbial mineralization results from adventitious precipitation of inorganic compounds led by their interactions with different metabolic processes and is defined as ‘biologically induced mineralization’.^3,4^ The most common minerals deposited by bacteria are calcium carbonates and calcium phosphates,^3^ and they usually accumulate on the surface of the bacteria (epicellular mineralization), which then become embedded in the growing crystals.^3,5^ Understanding the process of mineral deposition in bacterial biofilms is important for a number of reasons. On the one hand, pathological mineralization may lead to a wide range of diseases, including dental calculus^6^ that can be associated with serious conditions like periodontitis.^7^ Kidney stones are other examples of mineralized biofilms that form in physiological environments.^8^ On the other hand, bacterial biofilms are living materials with potential technical applications. Here, mineralization may serve as one possible strategy to modify their mechanical and other physical properties.^9,10^

Pathological calculi have been described from the mineral and microbiological points of view; yet, how mineralization is triggered is still unclear.^11^ In saliva, mineral precursors for hydroxyapatite (HA), octacalcium phosphate and whitlockite are present in supersaturated concentrations. These are the most frequent calcium phosphate crystal phases found in dental calculus.^12^ However, their concentrations are not high enough to cause spontaneous precipitation.^13^ Recently, alkaline phosphatase (ALP) has been identified as an indicator of periodontitis,^14^ which suggests that bacteria from the oral microbiome could exploit similar mechanisms as eukaryotic cells (e.g., bone cells) to induce mineral deposition in the mouth. Kidney stones have been historically linked to ion supersaturation in urine. While this remains considered as a risk factor,^15^ recent research showed that mineral precursor supersaturation is actually not more frequent in patients developing kidney stone disease than in control groups.^8,16,17^ This observation encouraged scientists to reconsider the role of bacteria in kidney stone formation where bacteria may not only become trapped, but could also play a significant role in stone growth.^8,18,19^

While the induction of calcium carbonate precipitation by *B. subtilis* and *E*.*coli* has been studied in the context of living and self-repairing materials,^20,21^ the present study will focus on calcium phosphates, which are also widespread and widely studied in the eukaryotic world.^22^ In mammalian bone, calcium phosphates are deposited in the form of HA through a finely tuned process that results in a functional material with high levels of spatial organization, complex morphologies and well-defined crystallo-chemical properties.^23^ This process is defined as ‘biologically controlled mineralization’, in contrast to the adventitious ‘biologically induced mineralization’ happening in most prokaryotes.^3^ Bone-specific ALP plays a key role in HA precipitation. It catalyzes the hydrolysis of monophosphates from organic sources (e.g., alkaloids, proteins) and makes them available for the interaction with cations (e.g., calcium).^11,24–26^ Bacterial ALPs are either found intracellularly, in the periplasmic space of gram-negative bacteria (e.g., *E. coli*)^27,28^ or extracellularly, either as secreted enzymes or bound to the cell surface (e.g., *S. aureus*).^29,30^ In *E. coli*, periplasmic ALP is encoded by the *phoA* gene.^27^ PhoA was shown to induce HA formation in the presence of individual *E. coli* bacteria;^31^ however, its role in biofilm formation remains to be investigated, as has recently been done for *S. aureus* biofilms.^24,32^

In this work, we used the *E. coli* K-12 strain W3110, a lab strain that expresses periplasmic PhoA (from now on referred as ALP),^33^ to grow biofilms on nutritive agar substrates supplemented with calcium chloride and β-glycerophosphate as a source of organic phosphate.^31^ The presence and distribution of minerals in the biofilms were detected via Calcein fluorescence. At smaller length scales, the minerals were localized with X-ray microtomography and with Focused Ion Beam with Scanning Electron Microscopy (FIB-SEM). Wide-angle X-ray scattering (WAXS) enabled to identify the mineral phase as HA. The crucial role of ALP in biofilm mineralization was finally demonstrated by the lack of biofilm mineralization on mineralizing medium supplemented with an ALP inhibitor, and the formation of mineral after depositing a droplet of bacterial ALP solution (without the bacteria) on the mineralizing medium.

## Results

### *E. coli* biofilms mineralize in presence of calcium and β-glycerophosphate

To determine the mineralization conditions of *E. coli* biofilms at the solid-air interface, we first used Luria Bertani (LB) agar substrates deprived of sodium chloride but supplemented with calcium ions and sodium β-glycerophosphate at different concentrations (Figure 1A). The β-glycerophosphate concentration was set to 10 mM. This value approaches the physiological concentration of phosphate in human saliva^12^ and is identical to the phosphate concentration at which *E. coli* can mineralize in liquid medium.^31^ Despite not being naturally present in mammalian saliva, β-glycerophosphate is traditionally used as a model for organic phosphates in dental plaque calcification.^34,35^ Two concentrations of calcium chloride were tested: a concentration of 1 mM corresponding to the typical ion concentration in human saliva^12,36^ (experiment named P-Ca 10:1) and a concentration of 10 mM, that has previously been used to induce mineralization in liquid medium^31^ (experiment named P-Ca 10:10). Finally, salt-free LB agar was used as a control condition (non-mineralizing medium). To localize mineral precipitation, Calcein Green (4 µM) was added to the nutritive medium as an indicator of calcium accumulation.^37^ We seeded the different agar substrates with suspensions of *E. coli* W3110 that constitutively express the fluorescent protein mCherry in their cytoplasm, and imaged the biofilms at different time points during 10 days of growth. Biofilms grew from the initial inoculation trace up to a diameter of about 18 - 19 mm within 10 days (Figure 1B). Biofilms grown on the medium with the lower Ca^2+^ concentration (P-Ca 10:1) did not show any relevant differences in mass and final size with respect to salt-free agar. Biofilms grown on the medium with the higher Ca^2+^ concentration (P-Ca 10:10) had a smaller final diameter and lower mass compared to the other two conditions. After 10 days of growth (Figure 1C), mCherry fluorescence indicated that most of the bacteria were concentrated in the center of the biofilm

**Figure 1.**
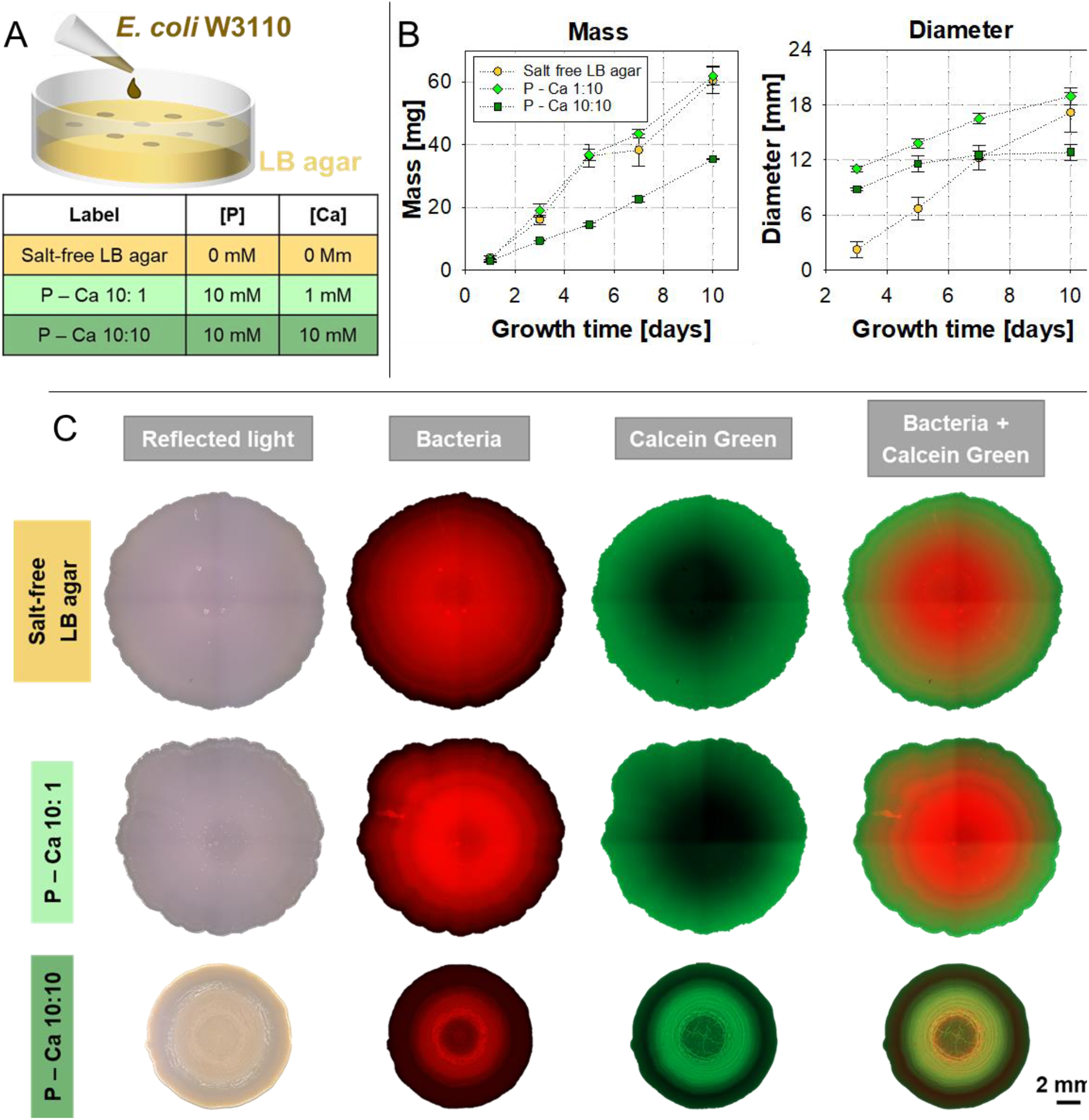
Detection of mineralization depending on different culturing conditions. A) Overview of the composition of the nutritive substrates with the respective Ca2+ ions and β-glycerophosphate concentrations. B) Diameter and wet mass of the biofilms, respectively estimated from the stereomicroscopy images and by gravimetry. C) Stereomicroscopy of biofilms after 10 days of growth in different conditions. Shown is the reflected light (first column), mCherry fluorescence to highlight the presence of bacteria (second column), Calcein fluorescence (third column) and overlaid mCherry and Calcein fluorescence (fourth column).

In contrast, Calcein fluorescence indicated high Ca^2+^ concentrations at the biofilm periphery. These different distribution patterns of bacteria and Ca^2+^ are consistent with the diffusion of Ca^2+^ and Calcein from the medium into the biofilm, while the intracellular (i.e. cytoplasmic) calcium concentration remains in the range from 100 to 300 nM.^38^ The emergence of these characteristic distributions is also shown in Figure S1, which reports fluorescent images taken at different times of growth (from day 3 to day 10). For biofilms grown in the P-Ca 10:10 condition, however, the Calcein signal was much brighter in the biofilm center and partially co-localized with the mCherry signal. This indicates a local accumulation of Ca^2+^ in regions with a high number of bacteria (Figure 1C and S1). Biofilms grown in these conditions also appeared opaque white, strongly suggesting that the P-Ca 10:10 growth condition favors calcium-based mineral precipitation in the biofilm.^37,39,40^

### Mineralization localizes close to the biofilm interfaces and involves both matrix and bacteria calcification

As fluorescence microscopy revealed non-uniform biofilm mineralization (Figure 1C), X-ray microtomography and FIB-SEM experiments were conducted to locate the minerals in the biofilm at different scales and in three dimensions (Figure 2). Microtomography was performed with fixed biofilms that were grown for 10 days on salt-free LB agar and on P-Ca 10:10 substrates. Biofilms grown on salt-free LB agar did not show a contrast difference with respect to the agar. On the contrary, P-Ca 10:10 biofilms clearly showed regions of higher gray levels close to the biofilm-air and biofilm-agar interfaces, which indicated a higher concentration of elements absorbing X-rays efficiently due to the high atomic number (like calcium). These regions were separated by a 50 µm thick layer within the biofilm that shows a lower degree of mineralization (Figure 2A).

**Figure 2.**
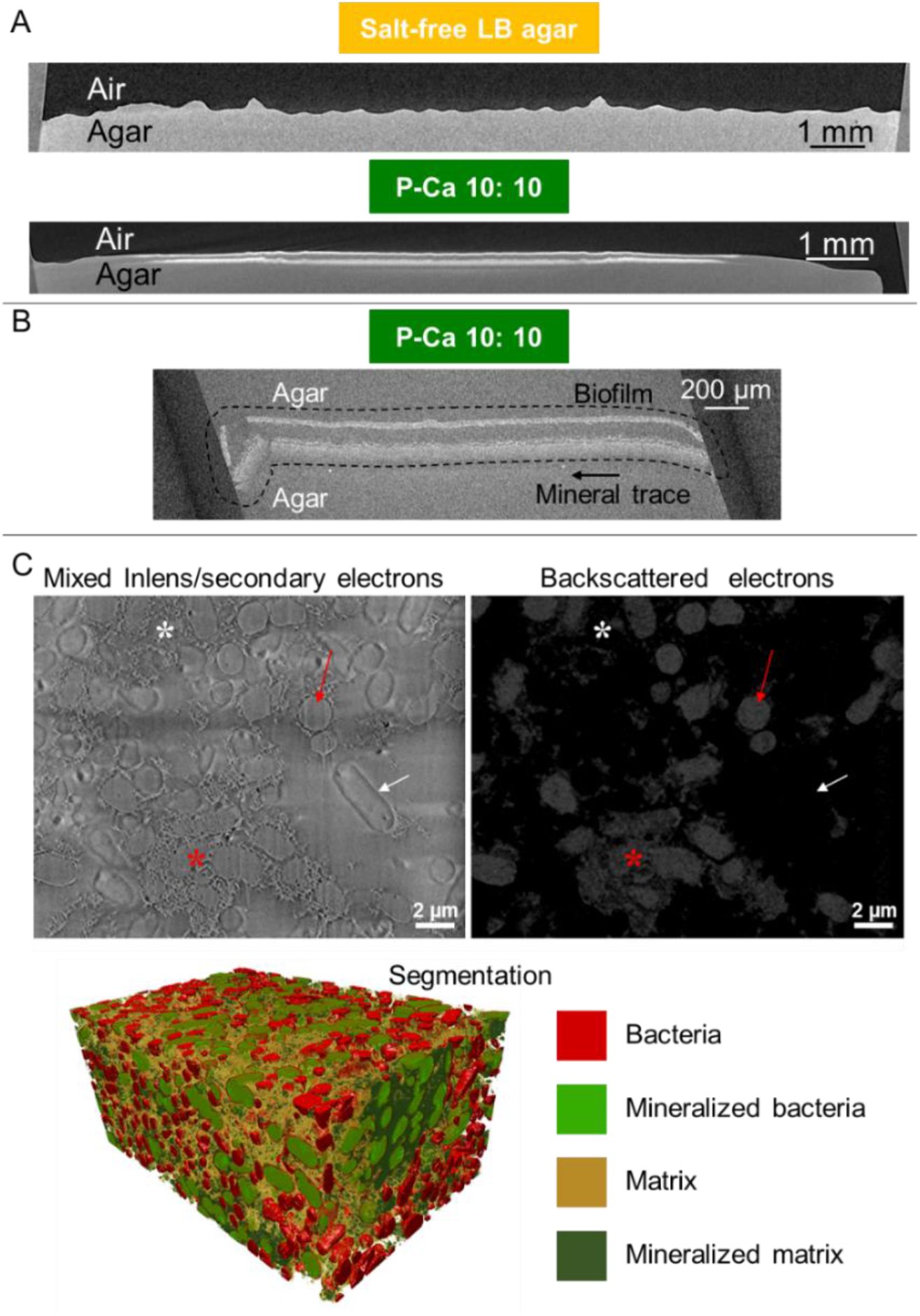
Mineral location at different length scale. A) Microtomography of biofilm cross-sections. Biofilms were grown on salt-free LB agar and P-Ca 10:10, and fixed prior to measurement. B) Microtomography in the central region (1.5 mm diameter) of a biofilm grown on P-Ca 10:10. The sample was stabilized by pouring another agar layer on top of the biofilm. C) FIB-SEM analysis of a biofilm grown on P-Ca 10:10. Top: mixed Inlens and secondary electron image, showing the sample topography (left) and backscattered electron image. The sample shows regions with different density (right). Arrows highlight mineralized (red) and unmineralized (white) bacteria, visible with the mixed Inlens/secondary electron detector. Only mineralized bacteria provide contrast with the backscattered electron detector. Stars show mineralized (red) and unmineralized (white) portions of the matrix. Bottom: Combining information from the backscattered and mixed Inlens/secondary electrons, it is possible to segment the sample and visualize all structures in 3D (see also video S2).

To achieve higher resolution, we prepared a biofilm sample of smaller volume. To stabilize the structure during sample preparation, a layer of agar was added on top of the biofilm. After solidification, a small cylinder (1.5 mm diameter; 2 mm in thickness) was punched out of the central mineralized region. Microtomography of this sample revealed that the mineralized layer at the biofilm-agar interface was thicker than the layer at the biofilm-air interface (Figure 2B). The brittle fractures of these two layers, which most probably occurred when punching out the sample, indicated a radical change of biofilm material properties (e.g., toughness) upon mineralization (Figure 2B, left side). These data also confirmed that the mineralized layer at the biofilm-air interface was not a measurement artifact due to the density mismatch at the air interface. Moreover, the higher resolution achieved for this sample revealed a light trace of higher gray levels within the agar substrate that correlated with the observation of white traces in the agar after removing the biofilms from the substrate for further processing (e.g., for weighing) (Figure 2B, black arrow). The presence of minerals in the agar substrate indicated that the biofilms are also able to induce changes leading to mineralization in their surroundings.

FIB-SEM provides resolution at the nanometer length scale and allows the visualization of individual bacteria and their surrounding matrix. After 10 days of growth in the P-Ca 10:10 conditions, samples of approximately 1.5 mm in diameter were punched out from the mineralized central region of a biofilm. FIB-SEM was used in the dual channel mode, allowing the simultaneous image acquisition from both a mixed Inlens/secondary electron detector as well as backscattered electron detector (Figure 2C). The mixed Inlens/secondary electron detector allowed the identification of rod-like bacteria as well as the surrounding fibrous matrix. The backscattered detector revealed bright mineralized regions while unmineralized areas appeared dark. Taken together, these images show that, in some cases, minerals completely encapsulated and filled the bacteria (Figure 2C, red arrows), while other bacteria remained completely unmineralized (Figure 2C, white arrows). The fibrous matrix wss easily distinguished in the mixed Inlens/secondary electron images. When viewed with the backscattered electron detector, also the matrix showed local differences in mineralization (Figure 2C). To distinguish the different regions in 3D (e.g., mineralized and unmineralized bacteria, mineralized and unmineralized matrix), the volume was segmented (Figure 2C, video S2). Video S2 shows some clusters of mineralized bacteria surrounded by mineralized matrix.

### Minerals in *E. coli* biofilms precipitate in the form of hydroxyapatite

X-ray powder diffractometry (XRD) was conducted on fixed and freeze-dried biofilms to confirm the presence of minerals and identify their phase. This preparation enabled a systematic and high-throughput analysis. We initially measured biofilms grown in control and P-Ca 10:10 conditions, using an in-house diffractometer (Bruker D8). Biofilms grown in P-Ca 10:10 conditions showed peaks at q-values around 18 nm^-1^ and 22.50 nm^-1^ (Figure S3A). Additionally, we analyzed the residual white stain in the agar, left by the mineralized biofilms after removing them from the surface (Figure S3B) as the microtomography analysis (Figure 2B) suggested mineralization in this region. Agar samples were prepared in the same way as biofilms and revealed peaks similar to those obtained in the mineralized biofilms. No scattering peaks were detected in salt-free LB agar and in P-Ca 10:10 agar sampled in a region that showed no white stain (Figure S3B).

More thorough analyses of the lyophilized biofilm powders were performed using synchrotron radiation in a wide-angle X-ray scattering (WAXS) setting (Figure 3). Biofilms cultivated on salt-free LB agar presented a broad single peak around q = 14 nm^-1^, similar to a freeze-dried overnight culture containing bacteria only (Figure 3A). Biofilms cultivated in presence of β-glycerophosphate and Ca^2+^ ions showed diffraction peaks typical for mineralization, depending on their respective growth stage. At low calcium concentration (P-Ca 10:1, Figure 3B), peaks at q = 22.33 nm^-1^ and at q = 32.38 nm^-1^ were visible only starting from day 10. The first peak can be traced back to the 211, 112 and 300 peak family of HA (Figure 3D). At high calcium concentration (P-Ca 10:10, Figure 3C), these peaks sharpened from day 5 to day 10, indicating crystal growth. Additionally, peaks around q = 18.10 nm^-1^ and 27.85 nm^-1^ were detected. These correspond to the 002 and 310 reflection planes of a reference HA powder (Sigma Aldrich), respectively (Figure 3C).

**Figure 3.**
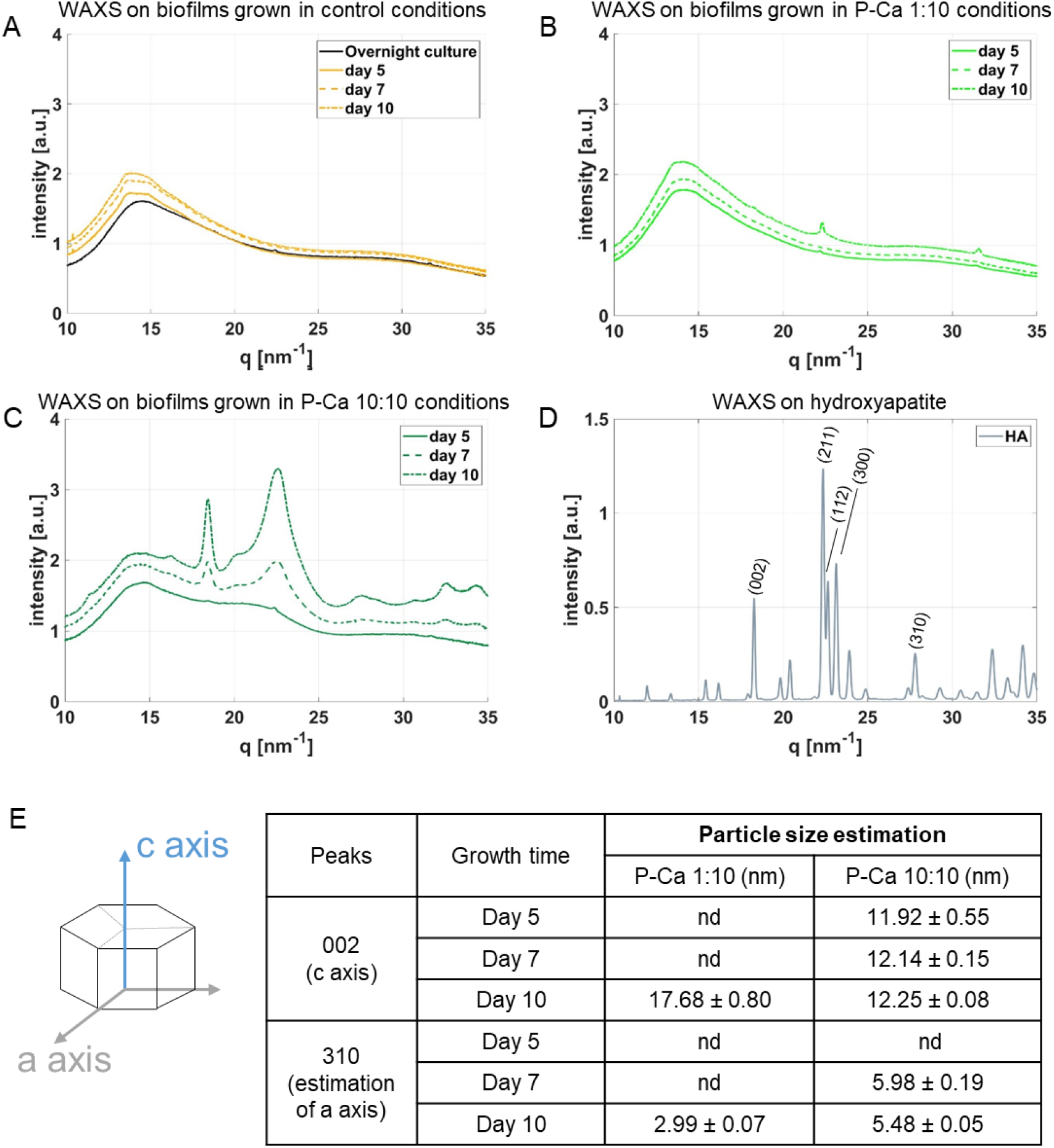
Mineral phase characterization through WAXS analysis. A-C) WAXS spectra of lyophilized biofilms after 5, 7 and 10 days of culture: A) biofilms grown on salt-free agar; B) biofilms grown on P-Ca 10:1 agar; C) biofilms grown on P-Ca 10:10 agar. D) Spectrum of HA as reference. E) Scheme of an HA crystal and estimation of HA particle size, using the 002 and 310 peaks (‘nd’ stands for ‘not detectable’). Data are shown as mean ± standard error of the mean (n = 12).

The Scherrer equation^41^ enables to estimate the crystal particle dimensions perpendicular to the corresponding planes (Figure 3E). Biofilms grown in P-Ca 10:10 conditions presented crystals with a dimension of about 12 nm along the c axis (peak 002, fig. 3E). Even though the average dimension increased over time, growth was not statistically significant (p > 0.01 under the hypothesis of normal distribution with a one-way Anova test, n = 12). The 310 peak allowed us to estimate the dimension of the diagonal of the hexagonal crystal base to range between 5.5 and 6 nm. Just as for the 211, 112 and 300 peak family, the 002 and 310 peaks were only detectable after 10 days when biofilms were grown in P-Ca 1:10 conditions. In these conditions, these peaks represented significantly longer, yet thinner particles of about 17.7 nm length and 3 nm base diagonal. The biofilm growth conditions thus seemed to influence the morphology of the resulting hydroxyapatite particles.

### ALP is required for hydroxyapatite deposition in *E. coli* biofilms

To test the hypothesis that mineralization is mediated by bacterial ALP, we implemented two strategies: (1) cultivating biofilms on a substrate containing the weak ALP inhibitor L-glutathione (IC_50_ ∼2 mM)^42^ and (2) testing the activity of the enzyme alone on P-Ca 10:10 agar plates.

The inhibitor L-glutathione^42^ was added in the P-Ca 10:10 agar and biofilms were grown under the same conditions as in the previous experiments. Comparing WAXS spectra of biofilms grown in P-Ca 10:10 conditions with and without inhibitor showed a delayed appearance of the HA peaks 002 and 310 in the presence of inhibitor. In the absence of inhibitor, peak 002 appeared very broad from day 5 and became sharper over time (Figure 4A, green lines). Peak 310 followed the same trend, yet started to be detectable from day 7. When the inhibitor was present (Figure 4A), both peaks 002 and 310 were less pronounced and only started appearing at day 7. WAXS thus confirms that mineral precipitation in the biofilms is delayed in the presence of a weak ALP inhibitor. Yet, a few HA crystals were still detectable in biofilms grown for 10 days with inhibitor. These crystals showed a c-axis of 13.05 ± 0.14 nm (mean ± standard error of the mean) and were thus significantly longer than the crystals found in biofilms grown without inhibitor.

**Figure 4.**
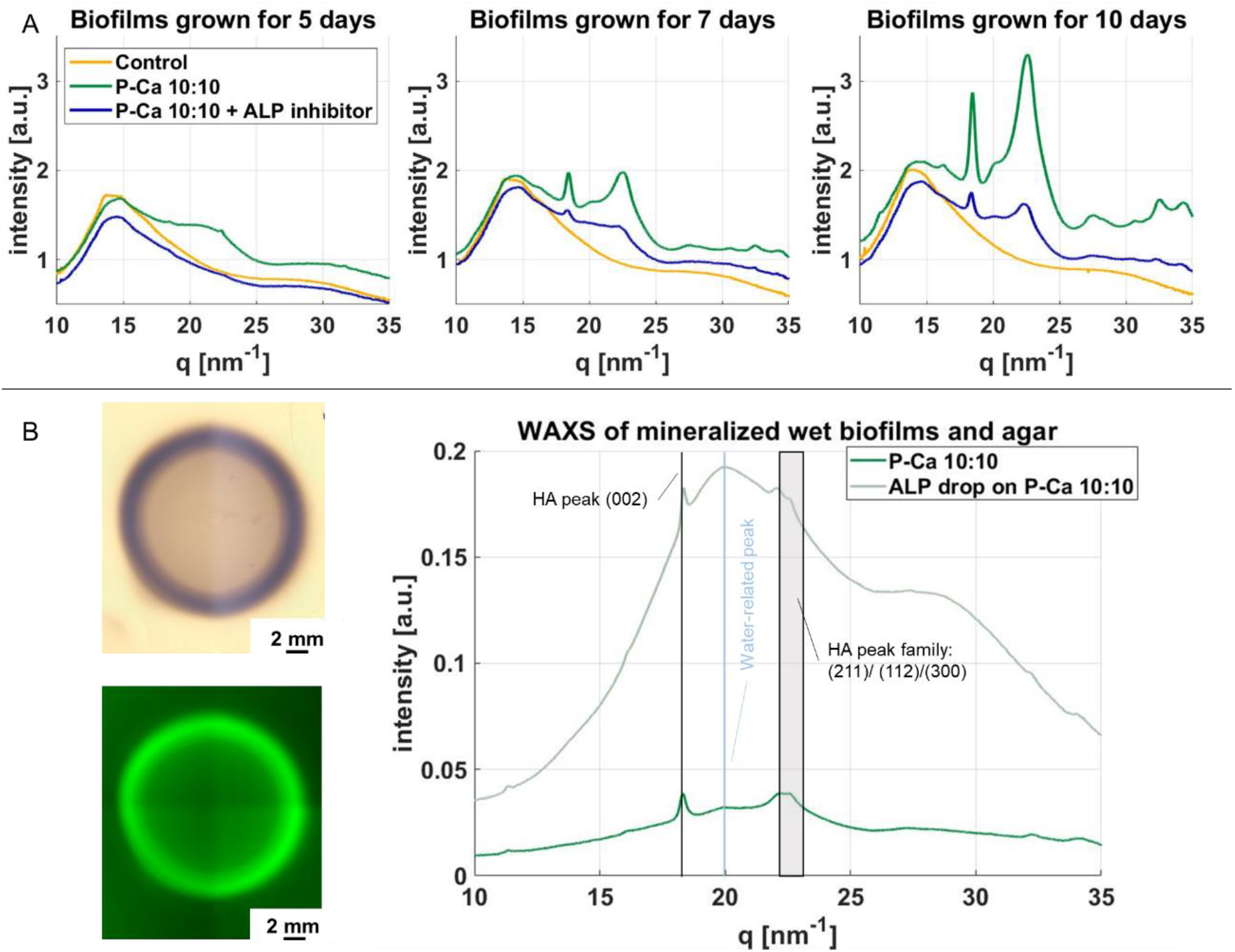
ALP role in biofilm mineralization. A) WAXS spectra of freeze-dried biofilms cultivated on LB agar, on P-Ca 10:10 agar and on P-Ca 10:10 agar supplemented with ALP inhibitor. B) Mineralization induced by an ALP solution spotted onto P-Ca 10:10 agar. Left: Stereomicroscopy showing reflected light (top) and Calcein fluorescence (bottom). Right: WAXS spectra of an area where a droplet of bacterial ALP solution was deposited on P-Ca 10:10 agar and a hydrated biofilm grown on P-Ca 10:10 agar.

To test ALP activity independent from the other biofilm components, a droplet of purified ALP was deposited on P-Ca 10:10 agar containing Calcein. After overnight incubation at 28 °C, the droplet left a marked dark trace with high Calcein fluorescence, indicating a higher local concentration of calcium (Figure 4B, left). Comparing the WAXS analysis of the hydrated material with a mineralized biofilm grown in P-Ca 10:10 conditions (not lyophilized) revealed that both spectra present the typical peaks at q = 18.1 nm^-1^, 22.5 nm^-1^ and (quite shallow) 27.85 nm^-1^ (Figure 4B). This shows that in the presence of calcium and β-glycerophosphate, ALP alone can induce the precipitation of HA minerals. Note that these spectra confirm that the minerals detected in the freeze-dried biofilms were not artifacts from sample preparation since they were also found in the native (wet) biofilms. These two independent experiments demonstrated that bacterial ALP is essential and sufficient in the biomineralization of *E*.*coli* biofilms grown in the presence of calcium ions and β-glycerophosphate.

## Discussion and Conclusion

This work shows that *E. coli* biofilms grown on a solid medium induce the precipitation of hydroxyapatite (HA) particles in the presence of calcium ions and an organic phosphate source. Our results also demonstrate that this biomineralization process is actively supported by bacterial alkaline phosphatase (ALP). Previous biogeological work has shown that individual *E*.*coli* bacteria can precipitate HA minerals when incubated in a solution of calcium ions and organic phosphates in absence of nutrients.^31^ This biomineralization process was particularly effective in *E. coli* cells overexpressing PhoA in the periplasm or in the cytoplasm. Most interestingly, extracellular mineral accumulation occurred independent of cellular PhoA localization, suggesting that either PhoA or inorganic phosphate is secreted into the extracellular space.^31^ Based on these observations, we implemented similar mineralization conditions in our *E. coli* biofilm growth protocol and tested a range of precursor concentrations that represent the range of human physiological conditions (Figure 1A). We showed that *E. coli* biofilm mineralization occurs in conditions where the concentrations of these precursors match those of the saliva (P-Ca 1:10),^12,36^ but with a time delay compared to conditions with higher concentrations (P-Ca 10:10). As human plasma contains on average 2.5 mM calcium,^43^ our experimental model is physiologically relevant and the P-Ca 10:10 condition can be representative of pathological situations where calcium ions tend to accumulate in given locations.^44^

The primary biological function of ALP is to mediate the availability of inorganic phosphate for metabolic processes. In prokaryotes, ALP is expressed when the concentration of inorganic phosphate is low, enabling the dephosphorylation of a wide range of organic phosphomonoesters, incl. phosphorylated proteins, lipids, carbohydrates and nucleic acids.^45^ While ALP plays an essential role in phosphate metabolism, its role in biofilm formation (e.g., matrix synthesis) and mineralization has not yet been investigated.^32^ Prokaryotic ALP dephosphorylates teichoic acid, a molecule that plays a key role for bacterial colonization on artificial surfaces. More importantly, dephosphorylation of organic phosphates and polyphosphates, liberates phosphate ions that may lead to supersaturation in the presence of Ca^2+^ ions. Purified bacterial ALP deposited on the mineralizing agar medium used for biofilm growth (Figure 4B) induced HA mineralization similar to the addition of the ALP to a solution of calcium ions and organic phosphates.^31^ This suggests a critical role of the enzyme in bacteria and biofilm mineralization, as confirmed by the absence of mineral formation when ALP inhibitor was added (Figure 4A).

Whether the formation of calcium phosphate minerals is favorable or not to the bacterial community is not clear and may depend on the physiological context. The FIB-SEM images show fully mineralized bacteria (Figure 2C). While mineralization causes the death of a large fraction of bacteria,^31^ the resulting composite material may offer protection to the small number of surviving bacteria and, overall, to the whole community. With this in mind, the less mineralized layer in between the mineralized top and bottom layers could become an environment where bacteria are protected from their surroundings (Figure 2A-B). The mineralized layers could block the diffusion of mineralization precursors and prevent mineralization of the entire biofilm and its bacterial population. From a population point of view, it could thus be worth the sacrifice of the mineralized bacteria. Such social behavior has already been described for bacteria communities in different contexts.^46,47^

There is no obvious correlation between the appearance of these layers and the cross-sectional structure of non-mineralized biofilms produced by the same bacteria.^48^ It is thus not clear if and how this layered geometry is collectively controlled by the bacteria or if it results from physico-chemical phenomena that involve a combination of diffusion and phase transitions. Indeed, layered periodic patterns of HA minerals, akin to Liesegang rings, spontaneously form in agar in presence of inorganic calcium and phosphate ions sources.^49^ Once phosphate ions are enzymatically released from β-glycerophosphate, our experimental conditions may resemble those of this study. This diffusion-based phenomenon may then explain the layered mineralization pattern in the biofilms and also the mineralization layer detected within the agar substrate, i.e. relatively far away from the bacteria (Figure 2B).

It remains an open question which key player is able to diffuse in the biofilm environment. Is ALP secreted by living bacteria or released by dead bacteria? Or is β-glycerophosphate taken up by the bacteria into the periplasmic space where inorganic phosphate is produced and subsequently secreted? Or is it even a combination of these possible processes? Further investigations are required to elucidate these aspects in order to clarify the mechanisms that determine the patterns of mineralization in biofilms. Similar to what is known about the role of collagen in bone apatite formation,^50^ biofilm matrix fibers may be involved in the nucleation of the mineral crystals due to favored interactions between ionic precursors and charged amino acids.^51^ In particular, curli fibers may serve as a scaffold to mediate HA mineralization as shown by *in vitro* studies with purified curli fibers.^52^ Moreover, the *E. coli* biofilm matrix may also influence the growth and morphology of the HA crystals, like the *B. subtilis* biofilm matrix was shown to influence the formation of calcium carbonate crystals.^53^

Altogether, the present work provides fundamental knowledge on biomineralization in microbial tissues as well as a platform to investigate biofilm calcification in a controlled environment that can be representative of physiological conditions. Insights into the interplay between metabolic state, biofilm structure and the accumulation of mineralization precursors will benefit not only the prevention of biofilm mineralization in the context of pathologies (dental calculus, kidney stones)^9,10,21^ but also to the engineering of biofilm-based living materials. Of particular interest are biofilm-based composites that are currently investigated as a potential source of sustainable materials with versatile properties adapted to various applications, such as self-repairing concrete and photosynthetic living materials.^20,54,55^

## Materials and methods

### Mineralization medium and biofilm inoculation

*E. coli* biofilms were cultivated at the solid-air interface, using tailored agar plates (15 cm diameter). Salt-free LB agar (Luria/Miller) plates (control medium) were prepared with 1.8% w/v of bacteriological grade agar-agar (Roth 2266), 1 % w/v tryptone (Roth 8952) and 0.5 % w/v yeast extract (Roth 2363). For the mineralizing media, calcium ions were added in the form of CaCl_2_ (Sigma Aldrich 223506). Sodium ß-glycerophosphate (Sigma Aldrich 35675) was added to the LB agar substrate as an organic source of phosphate. CaCl_2_ and sodium ß-glycerophosphate were sterile-filtered (0.22 µm) and added to the autoclaved and still liquid salt-free LB agar to reach a final concentration of 10 mM ß-glycerophosphate and 1 or 10 mM CaCl_2_. If calcium staining was required, a Calcein Green (Merck KGaa, 102315) stock solution (1 mM in 10 mM NaOH) was also added to the liquid LB agar to reach a final concentration of 4 µM. For the cultures with ALP inhibitor, a sterile-filtered L-glutathione (reduced, Sigma Aldrich G4251) solution was added to the liquid agar to a final concentration of 20 mM.

The *E*.*coli* K-12 strain W3110 was chosen as a well-characterized model strain.^33^ To visualize the bacterial cells using fluorescence stereomicroscopy, the bacteria were transformed with the plasmid pMP7604 (TetR; obtained from Guido V. Bloemberg, University of Zurich) that carries the gene for the fluorescent protein mCherry.^56^ The gene is under control of the strong *tac* promotor and the repressor was removed to allow for constitutive and strong expression of mCherry. It has been shown that this plasmid is stable and maintained in *E. coli* for at least 30 generations without selection pressure.^56^ This plasmid is thus an ideal marker for fluorescently labeling *E. coli* bacteria in a biofilm, avoiding the need for antibiotics.

In each plate, biofilms were inoculated with an array of 9 droplets, each using 5 µL of bacterial suspension (OD600 ∼5.0). The suspension was obtained from a single *E. coli* microcolony that was grown overnight in LB medium (Luria/Miller) (Roth X968) at 37 °C and 250 rpm. After seeding, the droplets were left to dry and the inoculated dishes were incubated at 28 °C for 5, 7 and 10 days. For the seeding on the dishes with the inhibitor, the LB medium for the overnight culture was also supplemented with 5 mM L-glutathione.

Apart from those used for the stereomicroscopy, all biofilms were fixed with 4 % paraformaldehyde (PFA, Bioster Ar106) in phosphate-buffered saline (PBS, Sigma-Aldrich P4417). The fixing solution was spread on the edge of each biofilm and left to react for 15 minutes. The excess solution was removed and substituted with fresh PFA. After another 15 minutes, the excess was again removed and the entire biofilm was covered with the fixing solution. The solution was left in contact with the biofilm for 2 hours. The excess was then removed and the biofilms were rinsed with the aforementioned PBS solution.

### Cryo FIB-SEM and Serial Surface View (SSV) imaging

Biofilms were grown for 10 days at 28 °C and fixed. Samples were collected from one complete biofilm, puncturing the central mineralized region at different locations with a disposable antistatic microspatula with a diameter of 2 mm (VWR international, Radnor, USA, PA). The samples were then sandwiched between two discs of 3 mm diameter, a type A and a type B gold-coated copper freezer hats (BALTIC preparation, Wetter, Germany), allowing an inner cavity of 0.5 mm in thickness with the addition of 10% dextran as cryo-protectant and cryo-immobilized in a HPM100 high pressure freezing machine (Leica Microsystems, Vienna, Austria). The sample carriers containing the samples were mounted on a cryo sample holder in the Leica EM VCM loading station (Leica Microsystems, Vienna, Austria) at liquid nitrogen temperature. The sample holder was then transferred using a VCT500 shuttle for freeze fracture and sputter coating (ACE600, Leica Microsystems, Vienna, Austria). After freeze fracture, the exposed samples were sputter coated with a platinum layer (thickness 8 nm). Finally, the samples were transferred to the Zeiss Crossbeam 540 microscope (Zeiss Microscopy GmbH, Oberkochen, Germany), using the VCT500 shuttle. Throughout the analysis, the biofilm samples were kept at a temperature below −145 °C.

FIB-SEM Serial Surface View Imaging was performed using a Zeiss Crossbeam 540 dual beam FIB-SEM (Zeiss Microscopy GmbH, Oberkochen, Germany). Three samples were used, resulting in three different stacks. The samples were elevated to a height of 5.1 mm, which corresponds to the coincident point of the two beams, and tilted to 54°. A trench of 20 μm length and 60 μm width was milled at 30 nA. The exposed surface was polished with a lower ion beam current (1.5 nA). The electron beam was then focused on the polished exposed tissue at 1.3 keV, 25 pA for stack 1 and 1.8keV and 50pA for stack 2 and 3. Images were sequentially collected using the “slice and view” protocol. The pixel size of the image prior to data collection was set at 8 nm for one stack and 10 nm for the two other stacks in the x and y directions. The slice thickness (z direction) was set to maintain an isometric voxel size, namely 8 nm for the first stack and 10 nm for the two others. All stacks were acquired with an image resolution of 2048 × 1536 pixels and in 8-bit grayscale. The first stack comprises 2082 slices, the second, 1340 slices and the third one, 1476 slices. For each stack, the dual channel option was enabled to simultaneously acquire images from the mixed Inlens/secondary electron (SE) detector and the backscattered electron (BSE) detector.

### Image processing

Stack alignment, image processing, segmentation and video generation were performed using Dragonfly software, Version 2021.1 (Object Research Systems (ORS) Inc, Montreal, Canada). BSE and mixed Inlens/SE images were first automatically aligned using the sum of square differences (SSD) method available in the slice registration panel. Curtaining artifacts produced by the beam during the milling process were corrected using the vertical destriping filter. A gradient domain fusion filter was applied only to the mixed Inlens/SE images in order to attenuate the discontinuities in the illumination from registered image stacks in the z direction. Finally, the contrast was improved and the noise reduced, using a convolution filter in all images. The segmentation was performed with the deep learning module in Dragonfly software. For training, 20 slices were used each data set, followed by a manual refinement using the ‘brush’ tool.

## Detection of calcium phosphate

### Fluorescence stereomicroscopy

After 5, 7 or 10 days of growth, pictures were taken from live biofilms with an AxioZoomV.16 fluorescence microscope (Zeiss, Germany). Images were taken in reflected light mode with an exposure time of 15 ms. The fluorescence signal from Calcein Green was collected with the 38 HE green fluorescent protein filter cube with the following specifications (excitation: 470/40 nm; beam splitter: FT 495 nm; emission 525/50 nm). The exposure time was 100 ms. mCherry fluorescence was detected using the 63 He red fluorescent protein filter cube (excitation: 572/25 nm; beam splitter: FT 590 nm; emission: 629/62 nm). The exposure time was 500 ms.

Biofilm diameters were estimated using custom-written MATLAB codes (Matlab 9.7.0 R2019b, MathWorks, Natick, MA). For the diameter calculation, the images taken in reflected light mode were converted to grayscale. With the function ‘edge’, we obtained skeletonized pictures and, after removing the noise with the function ‘bwareaopen’, we interpolated the remaining points with a circle and calculated the diameter of the circle.

### Microtomography

After 10 days of growth, fixed biofilms were cut out from the cultivation dishes using the upper part of disposable plastic Pasteur pipettes (12 mm diameter). The samples were kept sealed to avoid water evaporation. For the P-Ca 10:10 biofilms, a similar procedure was replicated in the center of the biofilms, but using disposable antistatic microspatulae with a diameter of 2 mm (VWR international, Radnor, USA, PA). Samples were imaged using an X-ray microtomography scanner (RX Solutions EasyTom160/150 tomographic unit) with a voxel size of 8 µm at 70 kV and 130 µA for the larger samples and a voxel size of 1.65 µm at 80 kV, 69 µA for the smaller samples.

### Identification of the calcium phosphate phase

To identify the mineral phase, we processed the biofilms to obtain a powder. After fixation, biofilms underwent freeze-drying in a lyophilizer (Lyo Alpha 1-2 LD; Martin Christ, Osterode am Harz, Germany) (pressure: 0.63 bar, temperature: −25 °C, overnight). Once freeze-dried, the powders were milled with an agate mortar, to obtain a finer powder.

### Wide angle X-ray scattering (WAXS)

Powdered samples were filled into a 2 mm Teflon sample holder with multiple holes of 5 mm diameter. The samples were overlaid with a Kapton film on both sides. Small-angle/wide-angle X-ray scattering (SAXS/WAXS) measurements were performed at the µSpot beamline at the BESSY II synchrotron (Helmholtz Zentrum für Materialien und Energie, Berlin, Germany).^57^ The complete spectra are shown in Figure S4. The measurements were carried out using a B4C/Mo Multilayer (2 nm period) monochromator and an energy of 15 keV. A sequence of pinholes was used to select a 30×30 µm^2^ spot size. Data were normalized over the primary beam intensity and the background was subtracted. Transmission through the sample was calculated from an X-ray fluorescence signal collected from a lead beam stop using a RAYSPEC Sirius SD-E65133-BE-INC detector. The detector was equipped with an 8 µm beryllium window, where the primary beam intensity was monitored using an ion chamber. Scattering data were collected with an Eiger 9M detector with 75×75 µm^2^ pixel area. For each sample, four spots were measured and diffraction patterns were collected with an exposure time of 15 seconds. Further data processing and reduction was done using the directly programmable data analysis kit (DPDAK).^58^ Diffraction patterns were radially integrated and the scattered intensity *I*(*q*) was calculated as a function of the momentum transfer *q*, defined as:

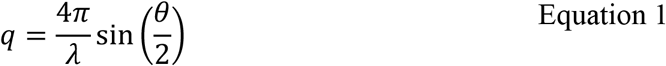

with *λ* and *θ* the photon wavelength and the scattering angle, respectively. The sample-to-detector distance was set to approximately 330 mm (0.1< *q* < 30 nm^-1^) and calibrated by using quartz powder. Data were analyzed with an in-house Python based software (Python 3.8.8).

### Mineralization in the presence of purified ALP

ALP (100U) from *E. coli* (Sigma Aldrich P5931, lot# 039M4019V) was dissolved in glycine buffer (0.1 M glycine/NaOH pH 10.4, 1 mM MgCl_2_, 1 mM ZnCl_2_). A droplet of this bacterial ALP solution (100 µL; 20 U/mL) was deposited on P-Ca 10:10 agar containing Calcein Green and incubated overnight at 28 °C. Samples were analyzed with stereomicroscopy and scattering experiments similarly to the biofilms.

## Supporting Information

The following files are available free of charge:

- Figure S1: Stereomicroscopy from 3 to 10 days of growth: reflected light and fluorescent signal
- Video S2: Segmented regions of the mineralized biofilm.
- Figure S3: Low resolution XRD spectra of Salt free-agar biofilms, P-Ca 10:10 biofilms and their traces on the agar
- Figure S4: Complete SAXS and WAXS spectra

## Data availability

Data and analysis tools are available upon request to the corresponding author.

## Author Contributions

The manuscript was written through contributions of all authors. All authors have given approval to the final version of the manuscript.

## Declarations

L.Z. gratefully acknowledges financial support from the German Research Foundation (Deutsche Forschungsgemeinschaft - DFG) for support through the DFG-Forschergruppe 2804 “InterDent”. Open access is funded by Max Planck Society. The authors declare no competing financial interest.

## Acknowledgements

We thank Dr. Guido V. Bloemberg (University of Zurich) for the gift of the pMP7604 plasmid and Prof. Regine Hengge (Humboldt-Universität zu Berlin) for kindly providing *E. coli* strain W3110. L.Z. acknowledges Christine Pilz for teaching her microbiological laboratory techniques. We thank Jeannette Steffen, Susann Weichold and Clemens Schmitt for their help with sample preparation. We also thank Daniel Werner for collecting the microtomography data and Dr. Chenghhao Li for the help provided calibrating the instruments at BESSY facilities. L.Z. acknowledge the research consortium of InterDent and the financial support of the German Research Foundation (Deutsche Forschungsgemeinschaft - DFG): DFG: FOR2804.

