## Supplementary Information for "Induced mineralization in *Escherichia coli* biofilms: the key role of bacterial alkaline phosphatase"

### Supplementary material

#### Stereomicroscopy from 3 to 10 days of growth

Stereomicroscopic pictures of the biofilms grown in control, P-Ca 10:1 and P-Ca 10:10 conditions. Pictures were taken with an AxioZoomV.16 fluorescence microscope (Zeiss, Germany) in reflected light mode with an exposure time of 15 ms. The fluorescence signal from Calcein Green was collected with the 38 HE green fluorescent protein filter cube with the following specifications (excitation: 470/40 nm; beam splitter: FT 495 nm; emission 525/50 nm). The exposure time was 100 ms. mCherry fluorescence was detected using the 63 He red fluorescent protein filter cube (excitation: 572/25 nm; beam splitter: FT 590 nm; emission: 629/62 nm). The exposure time was 500 ms.

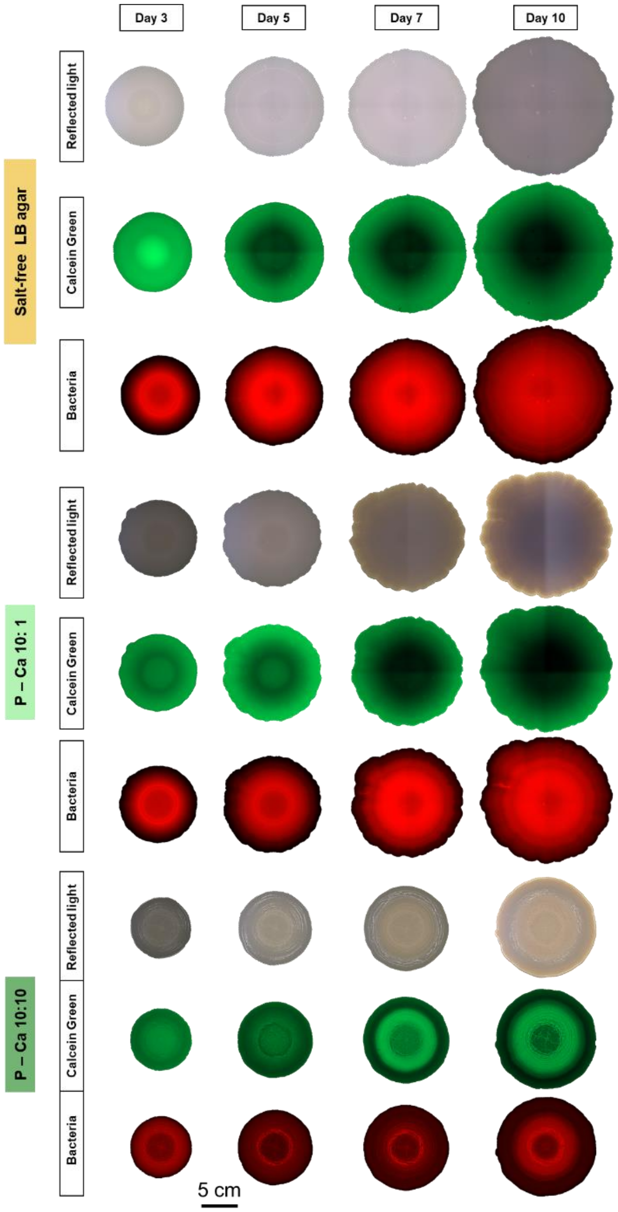


Figure S1. Stereomicroscopy from 3 to 10 days of growth: reflected light and fluorescent signal

#### Segmentation 3D reconstruction

Video S2 shows the segmented regions of the mineralized biofilm. It is possible to distinguish:

- Unmineralized bacteria (red)
- Mineralized bacteria (light green)
- Unmineralized matrix (light brown)
- Mineralized matrix (dark green)

#### Low resolution XRD spectra

XRD spectra were obtained with a X-ray diffractometer (Bruker D8) with the rotation of an X-ray copper tube and detector between 5° and 70°, with an increment between measurements of 0.02°/s. Exposure time was 1s for a total test duration of 54 minutes.

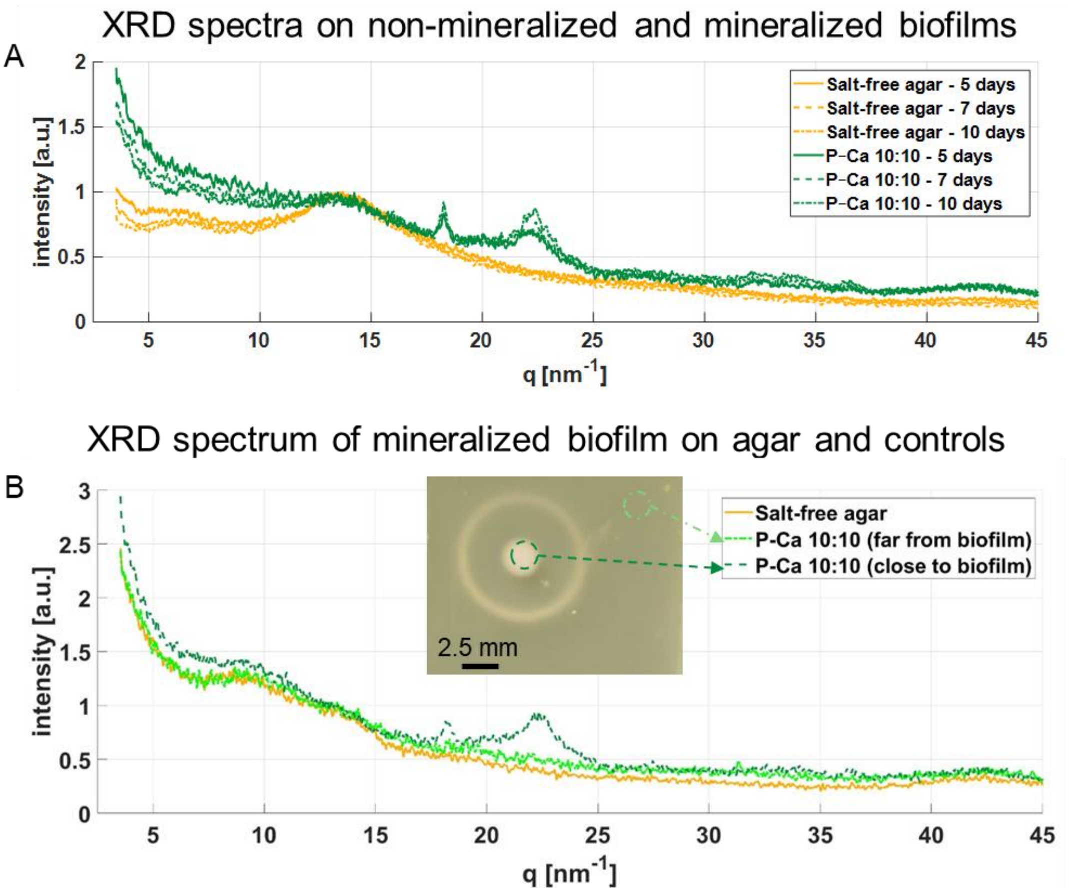


Figure S3. A) Preliminary XRD spectra on freeze-dried biofilms grown on salt-free LB agar and P-Ca 10:10 biofilms. B) XRD spectra of the trace left on the agar when growing biofilms on P-Ca 10:10 and respective controls (salt-free LB agar and P-Ca 10:10 agar sampled far from the biofilm).

#### SAXS and WAXS spectra

Complete spectrum from the small-angle/wide-angle X-ray scattering (SAXS/WAXS) measurements. Measurements were done according to the methods described in the Materials and Method section.

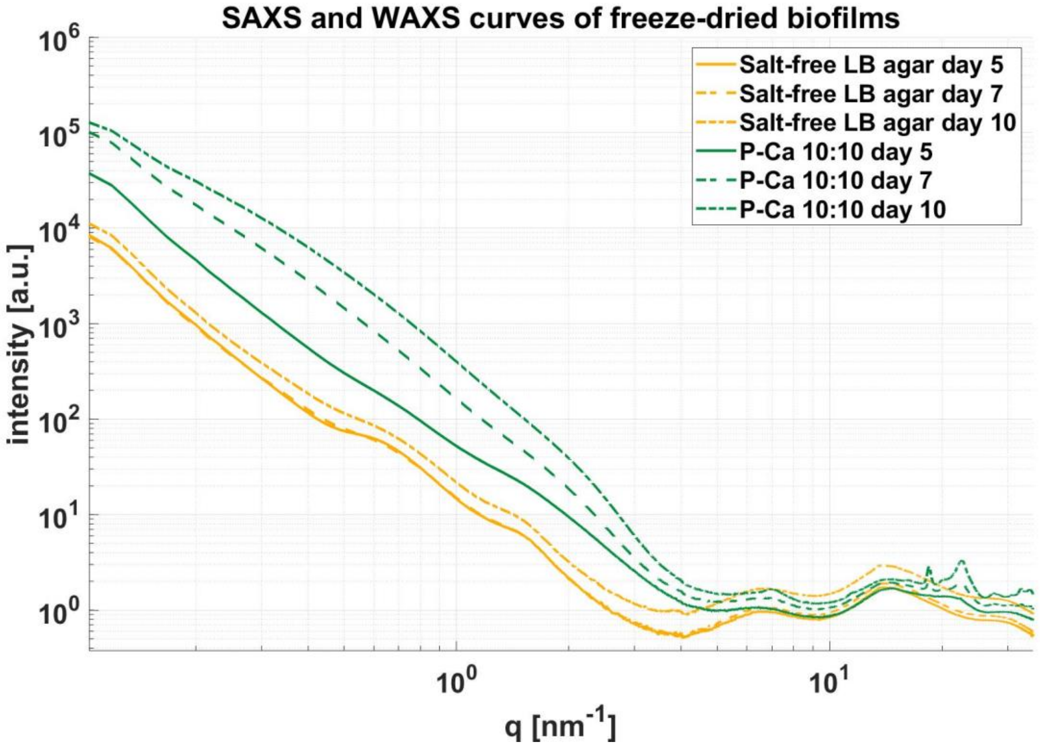


Figure S4. Complete SAXS and WAXS spectra.
